# Infected grapevines are poor hosts but can serve as source of pathogen transmission for SLF

**DOI:** 10.1101/2024.08.13.607612

**Authors:** Md Tariqul Islam, Crosley Kudla-Williams, Andrew D Harner, Michela Centinari, Cristina Rosa

**Affiliations:** Department of Plant Pathology and Environmental Microbiology, The Pennsylvania State University, University Park, PA, United States; Department of Plant Science, The Pennsylvania State University, University Park, PA, United States; Virginia Polytechnic Institute and State University, School of Plant and Environmental Sciences, AHS, Jr., AREC, Winchester, VA, United States

**Author notes:** Corresponding author: Md Tariqul Islam. Equal contribution (Md Tariqul Islam, Crosley Kudla-Williams).

**Keywords:** Spotted lanternfly, SLF, Pierce’s disease, *Xylella fastidiosa*, grapevines, etc

## Abstract

The potential of the invasive spotted lanternfly (*Lycorma delicatula* White; SLF) to serve as vector of plant pathogens is especially a concern for grapevine growers, as SLF are known to invade and can heavily infest vineyards, where the insects may encounter grapevines with multiple diseases. In this study, we have found that, when given the choice, SLF preferentially fed on healthy vines and that, when forced, feeding on Pierce’s disease (PD)-infected vines had negative effects on nymph development. Upon transmission trials, most of the recipient vines showed scorching symptoms typical of PD and one of the recipient vines resulted positive by qPCR. Our study suggests that SLF could be a vector of PD, however further experiments are needed to determine if transmission would occur under field conditions.

## Main manuscript

Spotted lanternfly (*Lycorma delicatula* White; SLF) is an invasive phloem-feeding insect native to Asia that is now established in the Eastern U.S. (Nixon et al., 2021). The first report of the SLF appearing in the U.S. was in 2014 in Berks County, Pennsylvania. Infestations have since been established across multiple states in the Northeast, Midwest, and South, including the states directly surrounding Pennsylvania, but also reaching as far west as Illinois and south to Tennessee and North Carolina. (N.Y. State Integr. Pest Manag., 2024). The rapid spread of SLF populations across the U.S. is a concern due to the threat the insect poses to agriculture. A voracious phloem-feeding insect, the SLF is known to feed on over 70 species of plants, including economically important crops, ornamentals, and hardwoods (Urban, 2020), but so far in the U.S. it has been a real problem on grapevines (Leach and Leach, 2020; Urban, 2020).

It is currently unknown if the SLF is capable of transmitting plant pathogens. Grapevines are susceptible to a wide variety of insect-transmitted diseases, and the introduction of a new vector could be devastating to growers. Hence, it is important to identify vectors and understand their role in pathogen spread to effectively manage and mitigate the impact of disease in vineyards. However, it is difficult to predict if an insect will be able to transmit certain pathogens without performing specific transmission tests, due to the wide diversity of insect vectors and a lack of characteristics known to be common to all vectors.

Among the most devastating vineyard diseases, Pierce’s disease (PD) has caused extensive economic damage in western and southern grape growing regions of the U.S., such as in California, Texas, and Virginia (Rapicavoli et al., 2018). This disease is caused by the xylem-limited bacterium *Xylella fastidiosa* subsp. *fastidiosa (Xff)*, which is transmitted, among other vectors, by *Homalodisca vitripennis*, or the glassy-winged sharpshooter (GWSS) (Rapicavoli et al., 2018), a robust xylem-feeding leafhopper in the same *Hemiptera* order as the SLF. Despite the xylem-limited nature of the pathogen, SLF may serve as a vector of PD because they could contact the xylem during their aggressive probing (Brooks et al., 2020). In addition, electropenetrography (EPG) studies have revealed that other phloem-feeding insects (Pompon et al., 2011), also belonging to the Fulgoridae family (Khan and Saxena, 1984), feed partially on xylem elements. Thus, there is a possibility that SLF can act as a vector of PD.

To address if the SLF can serve as vector, our study examined the effect of PD-infected grapevines on SLF development and determined if SLF is capable of transmitting PD.

From July to September 2022, we conducted experiments to assess the effect of PD-infected grapevines on SLF development and mortality. About 50 three-year old grapevines *Vitis vinifera cvs*. Cabernet Franc and Chardonnay, and two-year old Cabernet Sauvignon were grown in 19 L pots with custom substrate (2:1; Promix:topsoil, Sunshine® Perlite, Sun Gro Horticulture, Agawam, MA, Scotts® Premium Topsoil, The Scotts Company LLC., Marysville, OH) in a greenhouse at the Pennsylvania State University, University Park Campus, under APHIS permit. Vines were irrigated on alternate days, trained to one main shoot, and maintained to be approximately 1.5 m high. A group of Cabernet Franc vines in April 2022 and Sauvignon in 2022 and 2023 were inoculated with *Xylella fastidiosa* subs. *fastidiosa* strain Temecula 1, kindly provided by Dr. Roper, UCR, and grown at 28°C on solid PD3 medium for 7-10 according to Davis (Davis et al., 1981). A 20 µl drop of turbid cell suspension in pH 7 phosphate-buffered saline (PBS) was deposited onto the stem just above the third petiole from the base of the vine. The stem was pierced through the inoculum drop using a 20-gauge needle, so that the droplet was taken up into the xylem. The process was repeated on the opposite side of the stem to the initial droplet placement, for a total of 40 µl inoculum per vine (Purcell and Saunders, 1999). Control vines were challenged with PBS buffer alone. Vines were monitored for symptom development and tested to confirm infection.

Experiments were done to determine whether feeding on infected vines impacted the development and mortality of nymphs, as well as the feeding preference of the adults. To do so, fourth instar and adult SLF were manually collected from wild hosts across eastern Pennsylvania between July and September in 2022. Collected insects were transported to University Park and maintained until use in mesh insect cages (AgFabric, Wellco Industries, Inc., Coronoa, CA, USA) containing a potted grapevine or tree of heaven (*Ailanthus altissima* (Mill.).

We conducted SLF development and mortality experiments on 12 potted *Vitis vinifera cv. Cabernet franc* vines, 6 PD-infected and 6 uninfected, under greenhouse condition. Forty fourth instar SLF nymphs were added to each PD infected and uninfected vine, totaling 240 SLFs per treatment. Each vine was covered with a mesh insect bag modified with a zipper from Agdia (AgFabric, Wellco Industries, Inc., Coronoa, CA, USA) and arranged in a randomized complete block design with 6 blocks to account for potential variation in humidity and temperature within the greenhouse. SLF were counted and dead SLF were replaced on 6 dates throughout the experiment.

The development of 4^th^ instar nymphs to adult SLF was negatively influenced by PD infection (Figure 1A). A significantly higher number of nymphs developed into adults when left on healthy (uninfected) vines compared to PD-infected vines (*p =0*.*055*). Weekly SLF mortality was higher on PD-infected vines than on healthier vines (*p = 0*.*059*) only in the last week of the experiment (counted 46 days after infestation (DAI) (Figure 1B, left panel). When cumulative SLF deaths were considered, there was a significant increase of SLF death on PD-infected vines compared to healthy at three dates (39, 42, and 46 DAI; *p = 0*.*097, 0*.*041, and 0*.*028*, respectively, Figure 1B, right panel).

**Figure 1:**
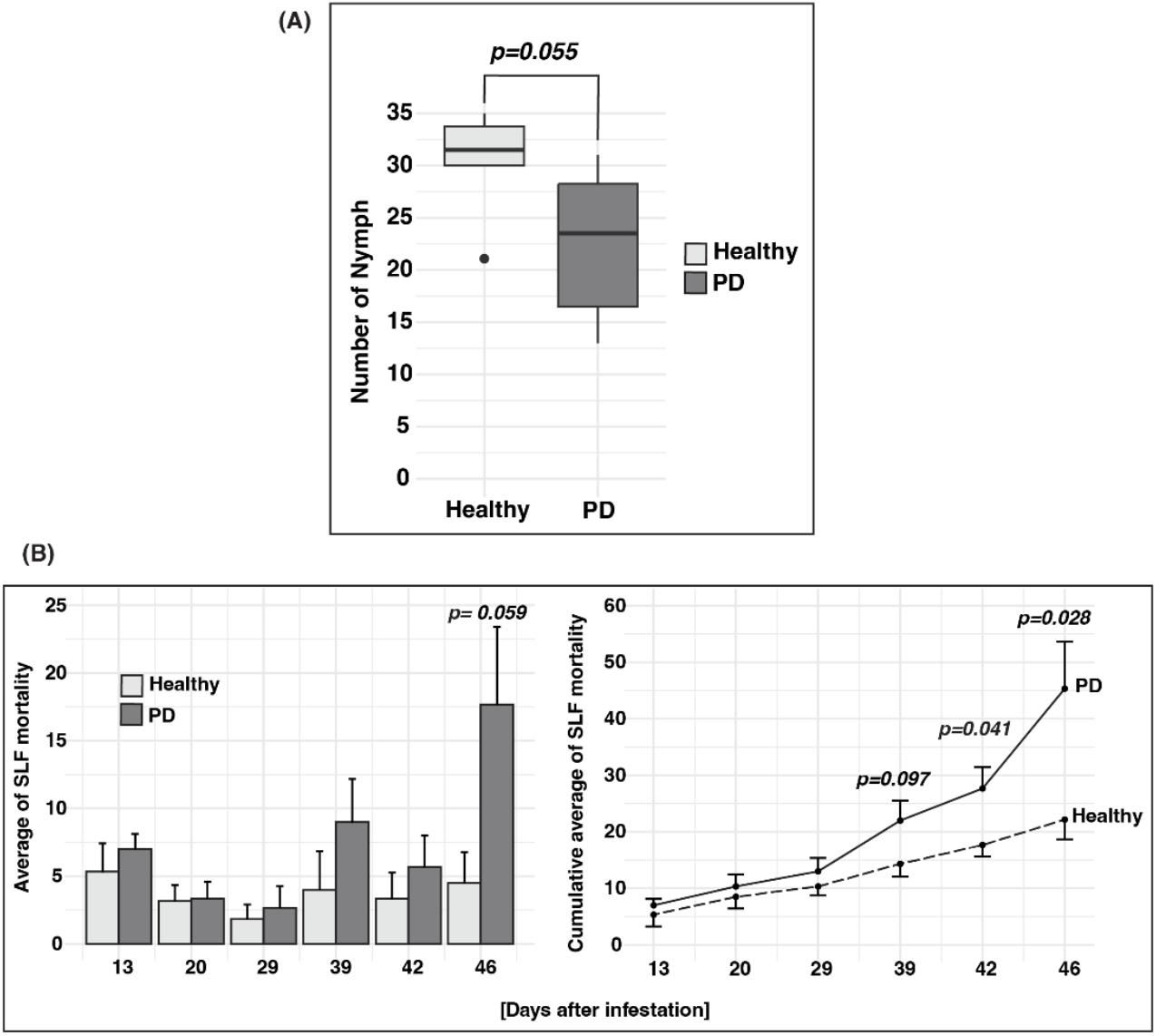
**Development of 4^th^ instar SLF nymphs to adults on healthy and PD-infected grapevines (A).** The box plots represent the number of SLF nymphs that molted into adults when caged on PD-infected (dark gray) or healthy vines (light gray). The box denotes the interquartile range of number of nymphs, the black bold line inside the box indicates the median and the whiskers represent maximum and minimum values. A total of 6 vines with 40 SLF per vine were used for each treatment (240 total SLF per treatment). Healthy vines displayed a higher number of successfully developed adult SLF compared to PD-infected vines according to one-way ANOVA with block as a random variable (*p=0*.*055*). **Mortality of SLF on healthy (uninfected) and PD-infected grapevines (B).** In the left panel, bars represent the mean ± SE number of SLF mortality counted approximately weekly after the infestation on healthy or PD-infected vines. A total of 6 vines were used per treatment, and adult SLF were added to the vines weekly to replace dead SLF and maintain 40 SLF per vine. When compared within each date, SLF exhibited a higher mortality on PD-infected vines at 46 days after infestation (*p=0*.*059*) according to one-way ANOVA. Whereas, in the right panel, lines represent the cumulative mean ± SE number of SLF mortality counted on the same days and it shows SLF has a higher cumulative mortality on PD-infected vines at 39, 42 and 46 days after infestation according to one-way ANOVA (*p=0*.*097, p=0*.*041, p=0*.*028*).

SLF feeding choice experiments were conducted from September to October 2022 using pairs of potted-grapevines: either two healthy *V. vinifera cv*. Chardonnay vines or one healthy and one PD-infected *V. vinifera cv*. Cabernet Franc vine. Paired vines were of similar age and growth stage and maintained under identical environmental conditions in the greenhouse. Each pair of vines had one shoot inserted into opposite ends of a tubular cage built using two tomato cages and covered with mesh (Figure 2A), with the shoots overlapping slightly in the middle.

**Figure 2:**
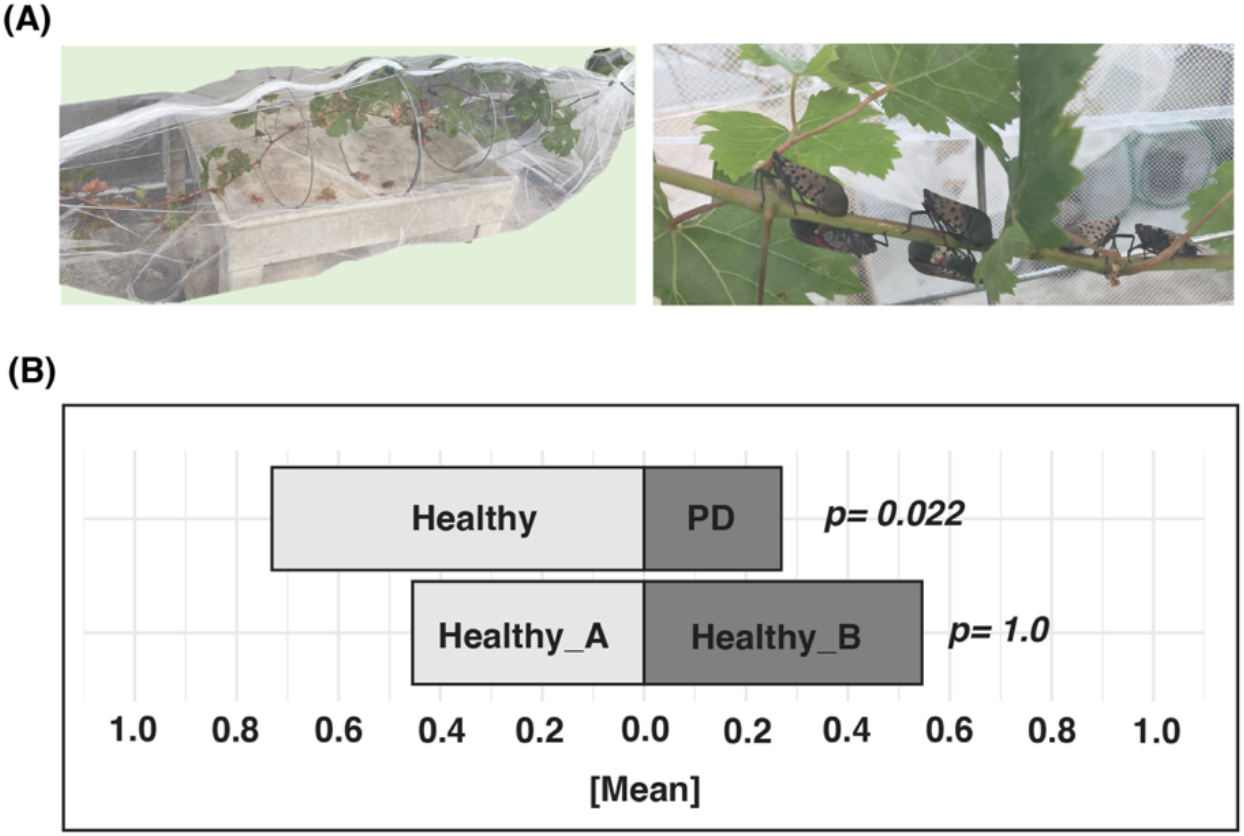
SLF choice experiment on PD-infected and healthy potted grapevines. **(A):** Image of the experimental setup and SLF feeding on the vines. **(B):** Bars represent the proportion of adult female SLF actively feeding on a given vine out of the total SLF actively feeding on all vines in the cage, averaged across 5-6 replicates (5 cages for HvsH and 6 cages for PDvsH, 10-12 SLF per cage). When given a choice between two healthy vines, SLF did not exhibit a preference according to according to a paired Wilcoxon signed-rank test (*p=1*.*0, we used* Wilcoxon signed-rank since the data was not normally distributed according to Shapiro test). When given a choice between a healthy and PD-infected vine, adult female SLFs exhibited a preference towards healthy vines according to a paired t-test (*p=0*.*022*).

For each biological replicate (defined as an individual cage with a choice test experiment in it), 10-12 female-adult SLFs or males were starved for one hour and then released into the center of the cage, equidistant from each shoot, in the morning or early afternoon. A total of 5 healthy vs. healthy and 6 PD vs. healthy choice tests were conducted. Observations occurred approximately every hour during daylight for two consecutive days. The number of SLFs on each vine or the cage were visually recorded for each observation as well as how many were actively feeding with stylet inserted into the plant. The number of dead SLFs was also noted at each observation.

Preliminary choice tests using two healthy vines revealed that adult female SLFs influenced feeding choices, with male SLFs following female SLF locations and feeding patterns within the cage. Consequently, only female SLFs were selected for the feeding preference experiments.

When given a choice between a PD-infected and a healthy vine, adult female SLFs preferred to feed on the healthy vine, as indicated by a significantly higher average proportion of SLFs actively feeding on the healthy vine compared to the PD-infected vine out of the total number of SLF in the cage (*p = 0*.*011*). The same result was obtained when considering the total number of SLF actively feeding on the vines, instead of the total number of insects in the cage (p = 0.022) (Figure 2B, top), indicating an overall preference for healthy vines over PD-infected ones.

As expected for controls in feeding experiments, when given a choice between two healthy vines, SLFs did not exhibit a preference for either vine in terms of the proportion of SLFs feeding on a given vine relative to the total SLFs in the cage (*p = 0*.*813*), nor relative to the total SLFs actively feeding throughout the experiment (*p = 1*.*0*) (Figure 2B, bottom).

We also conducted transmission assays to determine if SLF can vector *Xff* in grapevines. In 2022, 4^th^ instar and adult SLFs were used, while in 2023, only adult SLFs were used. Infected grapevines served as acquisition plants, and healthy grapevines as recipient plants, using *Vitis vinifera cvs*. Cabernet Franc (three-year-old) and Cabernet Sauvignon (two-year-old) in 2022 and 2023, respectively.

SLFs were given a 72-hour acquisition access period (AAP) on infected plants, followed by a two-week inoculation access period (IAP) on recipient vines. In 2022, four recipient vines were exposed to 60 nymphs each, and four to 40 adults each, beginning in July and August. In 2023, 6 vines were exposed to 33-50 adults each, starting in September. These numbers were chosen to mimic the natural infestations by SLF found in untreated vineyards and to maximize chances of survival up to the end of the transmissions, at which point only two-three SLF were still alive per each plant. August to September is also the time when SLF moves as adults into vineyards in Pennsylvania. Vines were monitored for symptom development and assayed for *Xff* four months post-IAP via qPCR. Dead SLFs were also collected at the end of the IAP in 2022, and during the AAP and one week into the IAP in 2023, then assayed for *Xff* by qPCR. Plant tissue for qPCR was prepared by combining 500 mg of petioles, midveins, and leaf blades from each plant. SLF samples were prepared by removing legs and wings from adults. Tissues were ground according to (Rowhani et al., 2000), and 2 µL of the prepared samples were used for qPCR. qPCR was performed using the Bio-Rad CFX96 Touch Real-Time PCR Detection System (Bio-Rad, Hercules, CA, USA). In 2022, qPCR was conducted using PerfeCTa SYBR^®^ Green FastMix (Quantabio, Beverly, MA, USA) as described by (Deyett et al., 2019) with thermocycling conditions: 95°C for 15 minutes, followed by 40 cycles of 55°C for 55 seconds, 72°C for 45 seconds, and 95°C for 15 seconds. In 2023, qPCR was conducted using SsoAdvanced Universal SYBR Green Supermix (Bio-Rad, Hercules, CA, USA) with conditions: 95°C for 30 seconds, followed by 39 cycles of 95°C for 5 seconds, and 60°C for 30 seconds. A dissociation protocol from 65°C to 95°C (5 seconds per cycle, 0.5°C ramp) was used to generate melt curves in both years. Primers used were XF-ITS-F6 (5’-GAG TAT GGT GAA TAT AAT TGT C-3’) and XF-ITS-R6 (5’-CAA CAT AAA CCC AAA CCT AT-3’) (Deyett et al., 2019).

We observed Pierce’s disease symptoms, including scorching of leaf margins, yellowing and reddening along leaf edges, defoliation of dried leaves, and shoot dieback near the growing tips, following nymph and adult transmission experiments in both years. Near complete defoliation and matchstick petioles occurred on some vines. Symptoms appeared approximately two months after SLFs were transferred from PD-infected to recipient vines, consistent with the timeline observed for *Xff* mechanical inoculations (Supplementary figure 1). In 2022, recipient vines were tested once with ELISA, twice with RPA, and multiple times with qPCR. However, they tested negative, whereas the positive controls (mechanically inoculated vines) were positive. In addition, samples taken from symptomatic tissues of recipient vines used in the Pierce’s disease transmission experiment were plated and yielded no colonies that were morphologically like colonies of known *Xff*. On the other hand, *Xff* was successfully isolated from grapevines inoculated mechanically with the bacteria, serving as a positive control and point of morphological comparison. These colonies grown from bacteria isolated from inoculated vines tested positive by qPCR, while the colonies from the experimental recipient vines tested negative. All plants, including the positive controls, were asymptomatic and negative for PD following vernalization in a walk-in cooler at 2-3 °C.

In 2023, adult SLFs left feeding on PD-infected grapevines and one transmission recipient grapevine tested positive for the bacteria by qPCR (Figure 3) ca. three months after transmission (January 2024), demonstrating that SLF likely ingests some xylem sap during feeding; and that PD transmission by SLF is possible.

**Figure 3:**
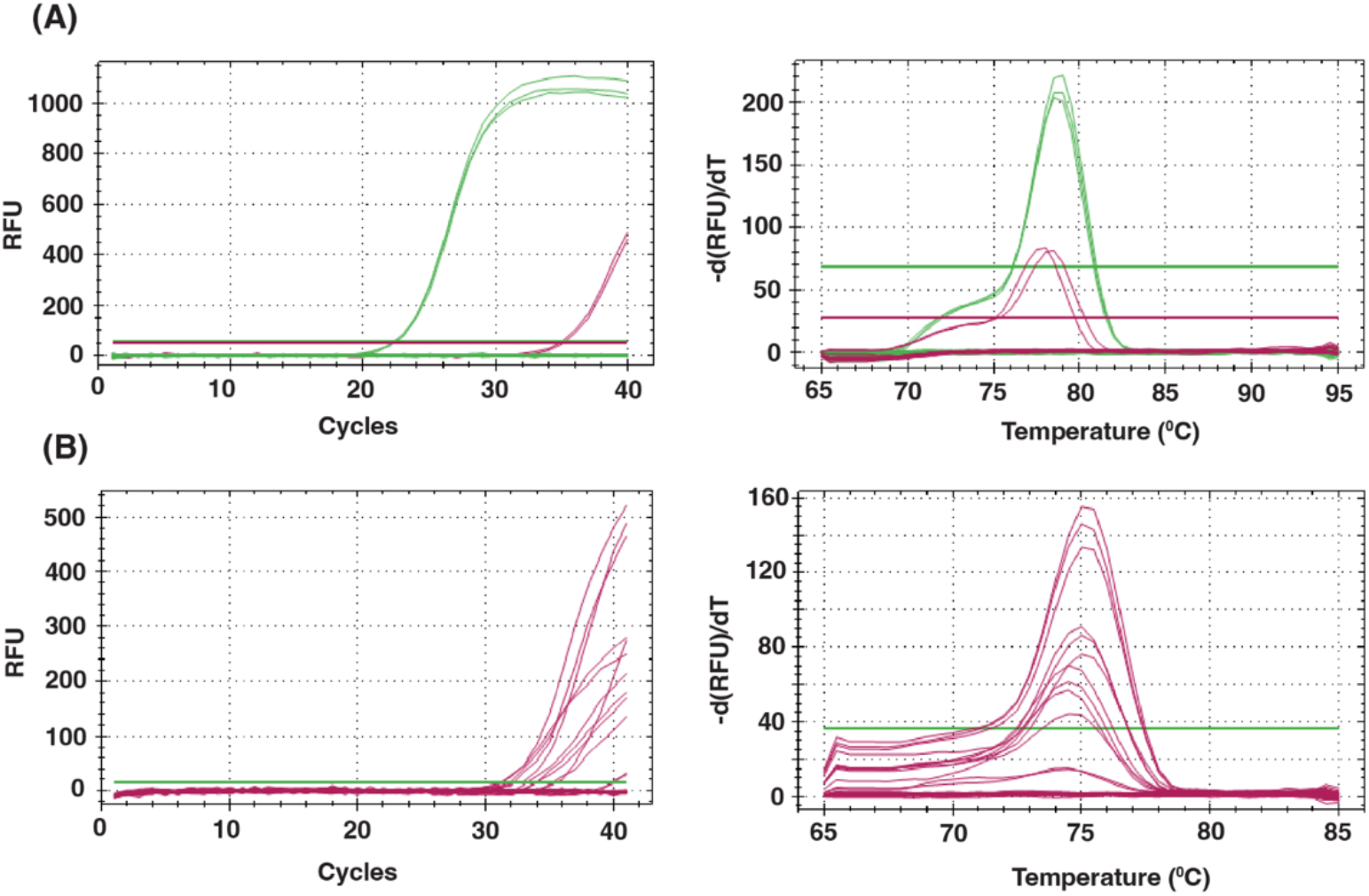
Transmission results by qPCR in 2023. The left panels show the Ct values for the amplification reactions, while the right the melt peaks. **(A):** qPCR data from grapevines. The green and burgundy lines indicate, respectively, the positive control grapevines used for PD acquisition and one of the 6 recipient vines. **(B):** qPCR data from dead SLF collected during the acquisition access period. We used leaf tissue samples from healthy vine as negative control, grape grinding buffer, glycine buffer, and water as no template controls.

While our data shows that SLFs may act as a vector of PD in greenhouse setup and that PD infected grapes seem to be nutritionally deficient for SLF and less palatable than uninfected plants, our experimental set up did not reflect the normal conditions for SLF in the field where the insects can freely move from vine to vine and the vines may have already been exposed to multiple diseases and be more or less attractive to SLF. Further experiments in the field are needed to understand the threat of SLF as PD vector.

## Acknowledgments

We want to thank Donald E Smith, Department of Plant Science for helping with the potting of vines.

## Reference

Brooks, R. K., Toland, A., Dechaine, A. C., McAvoy, T., and Salom, S. (2020). The inability of spotted lanternfly (Lycorma delicatula) to vector a plant pathogen between its preferred host, Ailanthus altissima, in a laboratory setting. Insects 11, 515.

Davis, M. J., French, W. J., and Schaad, N. W. (1981). Axenic culture of the bacteria associated with phony disease of peach and plum leaf scald. Curr. Microbiol. 6, 309–314.

Deyett, E., Pouzoulet, J., Yang, J.-I., Ashworth, V. E., Castro, C., Roper, M. C., et al. (2019). Assessment of Pierce’s disease susceptibility in Vitis vinifera cultivars with different pedigrees. Plant Pathol. 68, 1079–1087.

Khan, Z. R., and Saxena, R. C. (1984). Technique for demonstrating phloem or xylem feeding by leafhoppers (Homoptera: Cicadellidae) and planthoppers (Homoptera: Delphacidae) in rice plant. J. Econ. Entomol. 77, 550–552.

Leach, A., and Leach, H. (2020). Characterizing the spatial distributions of spotted lanternfly (Hemiptera: Fulgoridae) in Pennsylvania vineyards. Sci. Rep. 10, 1–9.

Nixon, L. J., Ludwick, D. C., and Leskey, T. C. (2021). Horizontal and vertical dispersal capacity and effects of fluorescent marking on Lycorma delicatula nymphs and adults. Entomol. Exp. Appl. 169, 219–226.

Pompon, J., Quiring, D., Goyer, C., Giordanengo, P., and Pelletier, Y. (2011). A phloem-sap feeder mixes phloem and xylem sap to regulate osmotic potential. J. Insect Physiol. 57, 1317–1322.

Purcell, A. H., and Saunders, S. R. (1999). Fate of Pierce’s disease strains of Xylella fastidiosa in common riparian plants in California. Plant Dis. 83, 825–830.

Rapicavoli, J., Ingel, B., Blanco-Ulate, B., Cantu, D., and Roper, C. (2018). Xylella fastidiosa: an examination of a re-emerging plant pathogen. Mol. Plant Pathol. 19, 786–800.

Rowhani, A., Biardi, L., Johnson, R., Saldarelli, P., Zhang, Y. P., and others (2000). Simplified sample preparation method and one-tube RT-PCR for grapevine viruses Presented at Meet. Int. Counc. Study Viruses Virus-Like Dis. Grapevine (ICVG), 13th, 12–17.

Urban, J. M. (2020). Perspective: shedding light on spotted lanternfly impacts in the USA. Pest Manag. Sci. 76, 10–17.

